# Autophagy degrades myelin proteins and is essential for maintaining CNS myelin homeostasis

**DOI:** 10.1101/2022.06.20.496817

**Authors:** Niki Ktena, Stefanos Ioannis Kaplanis, Irina Kolotuev, Alexandros Georgilis, Vasiliki Stavroulaki, Emmanouela Kallergi, Vassiliki Nikoletopoulou, Domna Karagogeos, Maria Savvaki

## Abstract

(Macro)autophagy comprises a major lysosome-dependent degradation mechanism which engulfs, removes and recycles unwanted cytoplasmic material, including damaged organelles and toxic protein aggregates. Although a few studies implicate autophagy in CNS demyelinating pathologies, its role, particularly in mature oligodendrocytes and CNS myelin, remains poorly studied. Here, using both pharmacological and genetic inhibition of the autophagic machinery, we provide evidence that autophagy is an essential mechanism for oligodendrocyte maturation *in vitro*. Our study reveals that two core myelin proteins, namely proteolipid protein (PLP) and myelin basic protein (MBP) are incorporated into autophagosomes in oligodendrocytes, resulting in their degradation. Furthermore, we ablated *atg5*, a core gene of the autophagic machinery, specifically in myelinating glial cells *in vivo* by tamoxifen administration (*plp-Cre*^*ERT2*^; *atg5*^*F/F*^) and showed that myelin maintenance is perturbed, leading to PLP accumulation. Significant morphological defects in myelin membrane such as decompaction accompanied with increased axonal degeneration are observed. As a result, the mice exhibit behavioral deficits. In summary, our data highlight that the maintenance of adult myelin homeostasis in the CNS requires the involvement of a fully functional autophagic machinery.

## Introduction

Myelin is the multilamellar, lipid-rich membrane that wraps the majority of vertebrate axons and ensures the rapid action potential propagation over long distances. Furthermore, this evolutionary innovation of vertebrates, serves as an insulator of the axon and participates in axon trophic support (Baumann & Pham-Dinh, 2001; Nave & Werner, 2014). In the central nervous system (CNS), myelin is produced by the membrane extension of specialized glial cells, the oligodendrocytes (OL). In CNS myelin, the most abundant proteins are the transmembrane proteolipid protein (PLP, 50%) and the small cytoplasmic myelin basic protein (MBP, 30%). Both are of utmost importance for myelin compaction with the former mediating compaction on the extracellular side of the membrane and the latter involved in myelin compaction between two cytoplasmic membrane leaflets. Genetic defects in myelin genes as well as disruption of myelin membrane in adulthood lead to severe motor and/or cognitive impairments in humans. Furthermore, myelin damage is linked to neurodegenerative diseases including multiple sclerosis, Alzheimer’s, Parkinson’s disease, etc. (Panfoli *et al*, 2014; Joseph & DeLuca, 2016).

Recent work has contributed to the notion that myelin is not a static structure, as initially considered, but it undergoes constant turnover throughout life to retain its plasticity (O’Rourke *et al*, 2014; Sampaio-Baptista & Johansen-Berg, 2017). It has been shown that there is a constant replenishment and degradation of myelin constituents in order to avoid functional decline of the membrane (Peters, 2002; Bartzokis, 2004; Young *et al*, 2013). The question that arises is by which mechanism myelin and myelin proteins are degraded during this remodeling. A very recent study pointed to the role of microglia that phagocytose myelin sheaths during developmental myelination (Huges & Appel, 2020), although microvascular endothelial cells and astrocytes have also been previously reported to engulf myelin debris (Ponath *et al*, 2017; Zhou *et al*, 2019).

Macroautophagy (hereafter referred to as autophagy) comprises an evolutionarily conserved pathway delivering proteins and damaged organelles to lysosomes for degradation (Ariosa & Klionsky, 2016). It initiates with the engulfment of intracellular cargo by the phagophore, which expands to form a double-membrane vesicle, termed the autophagosome. The whole process is very well orchestrated by a network of autophagy related genes (ATGs), such as the ATG12 conjugation system (Atg12-Atg5-Atg16), that promotes the formation of MAP1LC3/LC3 (microtubule associated protein 1 light chain 3)-positive phagophores (Rubinsztein, 2015; Dikic & Elazar, 2018). Following this step and through a series of conjugation events, the autophagosome will finally fuse with the lysosome to form a structure called autolysosome, which constitutes the final step of autophagy and is responsible for the degradation (Mizushima, 2011; Dikic & Elazar, 2018).

Though initially perceived as a process in bulk, emerging evidence clearly suggests that selective autophagy can also serve specific functions and degrade diverse substrates, including lipid droplets (lipophagy), aggregated proteins (aggrephagy), mitochondria (mitophagy) etc (Kirkin & Rogov, 2019). Recently, a new form of selective autophagy was described for the degradation of myelin surrounding peripheral nerves after injury and attributed to Schwann cells (myelinophagy; Gomez-Sanchez *et al*, 2015). The authors showed that activation of autophagy is an early and essential step for myelin clearance after trauma and subsequent axonal regeneration. Similar studies support the Schwann cell autophagy-mediated myelin clearance (Jang *et al*, 2015; Weiss *et al*, 2016; Brosius Lutz *et al*, 2017). In contrast to peripheral nervous system (PNS) and Schwann cells, in CNS and OLs the role of autophagy is not completely understood. Recent work demonstrated that autophagy in OLs is involved in both CNS injury and recovery (Munoz-Galdeano *et al*, 2018; Saraswat Ohri *et al*, 2018). Moreover, recently it was suggested that autophagy plays a key role in the survival and differentiation of OL progenitor cells and proper myelin development (Bankston *et al*, 2019).

The aim of this study was to elucidate the role of autophagy in mature OLs and in CNS myelin homeostasis. Our findings strongly support autophagy as an essential mechanism for OL maturation and maintenance during adulthood. More specifically, we demonstrate for the first time that autophagy plays a crucial role in maintaining cellular homeostasis in adult CNS as a basic degradative mechanism for myelin proteins and this mechanism is supported by our *in vitro* data as well. Upon ablation *in vivo*, myelin proteins accumulate with severe morphological and behavioral defects in mice. Taken together, these results provide novel insights into the mechanism by which CNS myelin itself can be degraded, other than via microglia and astrocytes as has been established already.

## Results

### Inhibition of autophagy results in maturation defects in oligodendrocytes

In an effort to understand the importance of the autophagy pathway in CNS myelin, we first tested its effects in the maturation of OLs. For this reason, we established primary OL cultures derived from newborn P2 C57BL/6 mice, where MBP+ OLs were analyzed. MBP+ mature OLs can be categorized into five morphological stages (stages 0-4), according to their maturation index. Specifically, according to the literature (Lee *et al*, 2020), we identified: stage 0-1 as cells with primary or multiple processes and without myelin membranes, stage 2-3 cells with increased ramification branching and the initiation or extension of myelin sheet formation and finally, stage 4 OLs, which represent cells at their final maturation that display a fully developed myelin membrane sheet surrounding all the cell branches (Figure 1A).

**Figure 1:**
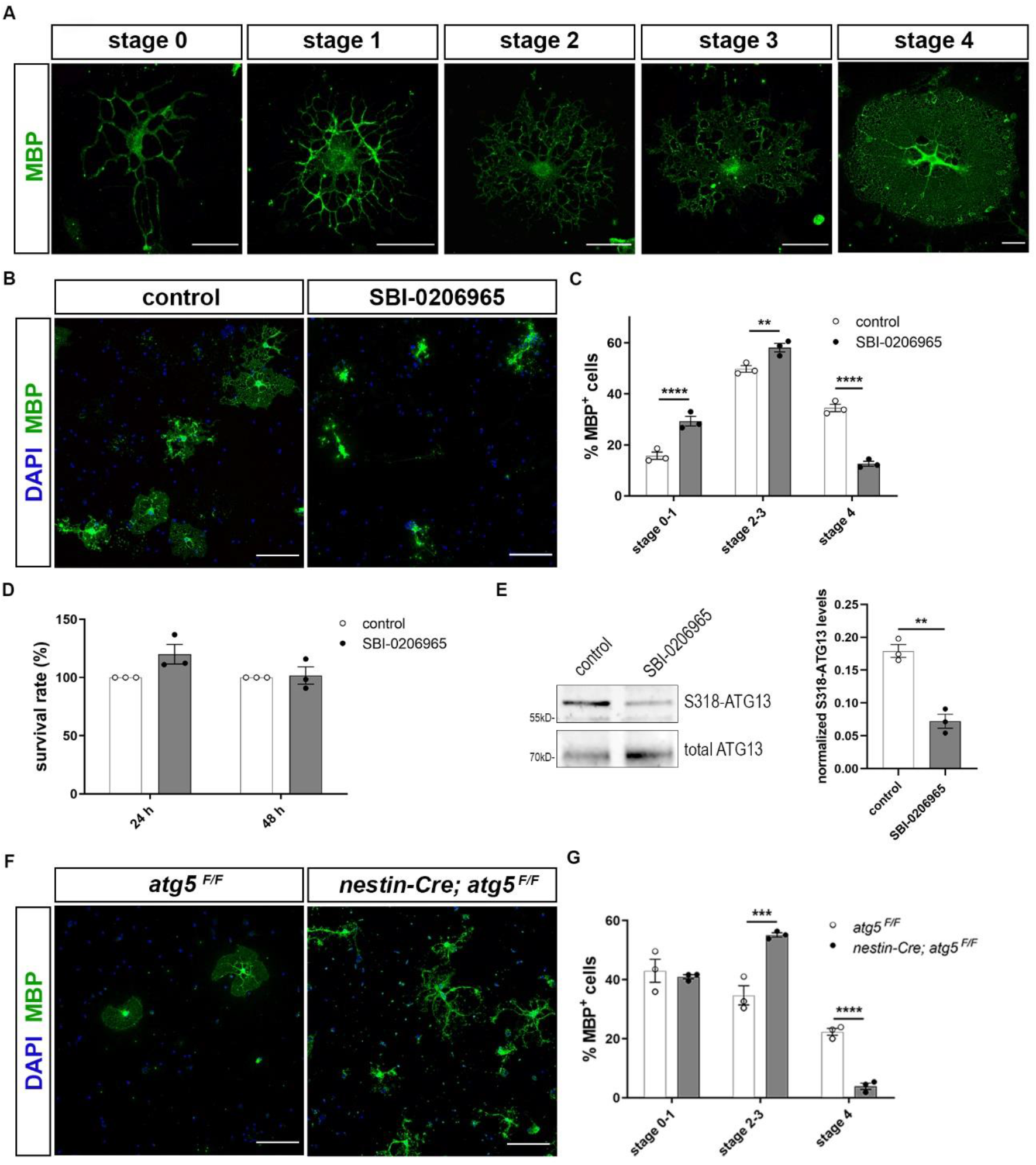
Pharmacological and genetic inhibition of autophagic vesicle biogenesis results in maturation defects in oligodendrocytes. A. Representative examples of mature MBP-expressing OLs (MBP in green) that can be subdivided in five maturation stages. Scale bars: 30 μm. B. Confocal images of DIV5 primary oligodendrocytes (OLs), immunostained for MBP (green) and DAPI (blue). Cells were either vehicle treated (control), or treated for 5 days with 1μM SBI-0206965. Scale bars: 100 μm. C. Percentage of the MBP+ OLs found in the categories described above. For the quantification of OLs complexity, the Two-way ANOVA (Sidak’s multiple comparisons test) was used. D. Western blot analysis and quantification of DIV5 primary OLs with an antibody against S318-ATG13, and normalized for total-ATG13. Cells were either vehicle treated (control) or treated with SBI-0206965 (1μ?) for 3 hr. Student’s t-test was used to determine statistical significance. E. Cell viability analysis by MTT assay performed on DIV5 OLs that were either vehicle treated (control) or treated with SBI-0206965 (1μ?) for 24 h or 48 h. Student’s t-test was used to determine statistical significance. F. Representative confocal images of DIV2 primary control (*atg5* ^*F/F*^) and cKO (*nestin-Cre; atg5*^*F/F*^) immunostained against MBP (green) and DAPI (blue). Scale bars: 100 μm. G. Quantification of the percentage of the MBP+ OLs found in the categories described above. For the quantification of OLs complexity, the Two-way ANOVA (Sidak’s multiple comparisons test) was used. Data information: Data are shown as mean ± SEM. Three independent experiments were used in each case. **p≤ 0.01, ***p≤ 0.001, ****p≤ 0.0001.

We initially performed pharmacological blockage of autophagy, using a potent and selective inhibitor of the serine/threonine autophagy-initiating kinases ULK1, named SBI-0206965 (Egan *et al*, 2015; Figure 1B). After 5DIV, these cultures were immunostained for MBP and DAPI and observed with confocal microscopy. Upon pharmacological blockage of autophagosomal formation, we noticed maturation defects in SBI-treated cells, since there were fewer OLs reaching final maturation stage 4 (Figure 1C). In parallel, we detected an increase in the proportion of early stage OLs (stage 0-1 and 2-3 OLs) in SBI-treated cultures. In order to ensure that SBI-0206965 does not affect cell viability in these cultures, we performed the MTT assay (Figure 1D). SBI-0206965 administration has no toxic effects on cell viability as observed. Moreover, protein levels of S318-ATG13 (normalized over the total ATG13 levels), a downstream target of ULK1 kinase, were found decreased in the case of SBI-0206965 treatment, as expected (Figure 1E).

To further validate the importance of autophagy in OL maturation, we decided to examine primary OLs from mice in which autophagy is genetically ablated. To this end, we performed primary OL cultures from P2 *nestinCre; atg5* ^*F/F*^ mice (Hara *et al*, 2006) and littermate controls (Figure 1F). Nestin promoter drives Cre recombinase expression in neuronal and glial cell precursors. Similar to the pharmacological blockage, genetic ablation of *atg5* in pure oligodendrocytic cultures demonstrated maturation defects in mutant cells (Figure 1G).

### Autophagy depletion leads to PLP accumulation *in vivo*

We further investigated the role of autophagy in OLs in an *in vivo* model. To this end, and in order to study the contribution of autophagy in mature OLs specifically, we crossbred the *atg5* ^*F/F*^ mice with the tamoxifen-inducible *plpCre* ^*ERT2*^ mice in order to eliminate the core autophagic component *atg5* exclusively in myelinating glial cells after tamoxifen administration. We first verified the successful recombination and the pattern of Cre activity by crossing our line with a transgenic reporter mouse line, namely mT/mG (Muzumdar *et al*, 2007). In this line, tdTomato (mT) fluorescence expression is widespread in cells, while the red fluorescence is replaced by the cell membrane-localized EGFP (mG), in Cre recombinase-expressing cells. For the tamoxifen injections, we followed the same experimental design used in our autophagy depleted mice *(plpCre* ^*ERT2*^; *atg5* ^*F/F*^), thus starting injecting tamoxifen into double-transgenic offspring (arising from *mT/mG*; *plpCre* ^*ERT2*^ breeding) at the age of 2.5 mo and analyzing these animals 3 months later, at 6 mo (Figure S1A). As shown in confocal images of both optic nerve sections (Figure S1B) and sagittal brain sections (Figure S1C) of tamoxifen injected *mT/mG*; *plpCre*^*ERT2*^-and *mT/mG*; *plpCre*^*ERT2*^+ mice, GFP expression is present strictly in myelin tracts of the latter, and absent in the *mT/mG*; *plpCre*^*ERT2*^-tamoxifen injected littermates. After verifying the efficiency of the recombination, we injected our mice (*plpCre* ^*ERT2*^; *atg5* ^*F/F*^) following the same tamoxifen administration protocol at 2.5mo and performed analysis on 6mo control (*atg5* ^*F/F*^) and cKO (*plpCre* ^*ERT2*^*+*; *atg5* ^*F/F*^) animals. Mice of both genotypes were treated with tamoxifen. The sufficiency of autophagic depletion in cKO mice was further validated via western blot by increased p62 levels (Figure 2A).

**Figure 2:**
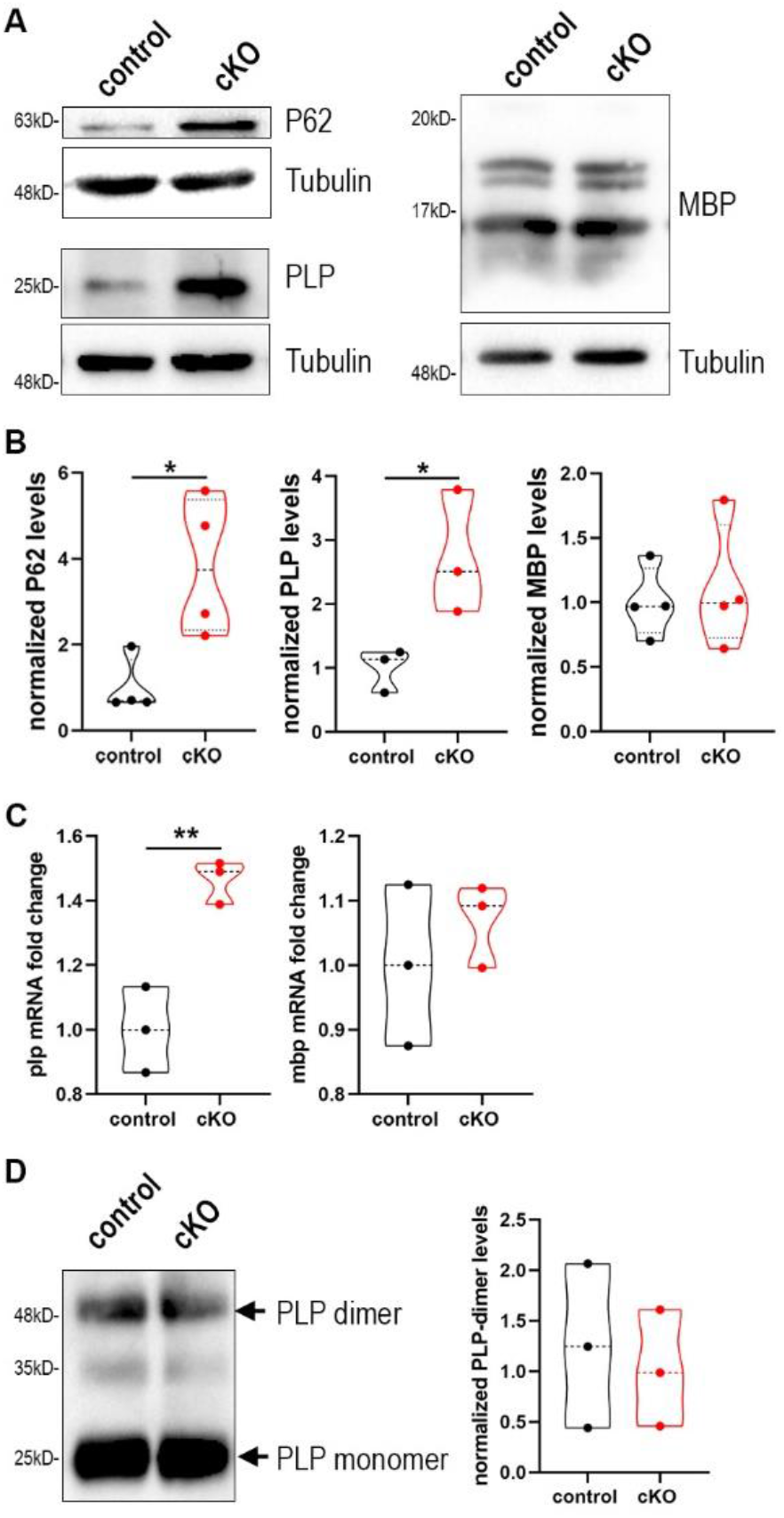
Blockage of autophagy leads to increased plp mRNA and protein levels. A. Western blot analysis with antibodies against P62, PLP and MBP proteins in optic nerve lysates of 6mo control and cKO mice. Representative images are depicted. B. Quantification of normalized protein levels. C. Graph showing quantification of quantitative real-time PCR in mouse forebrains of 6mo control and cKO mice. D. Western blot analysis, with antibody against PLP, of native membrane extracts (NME) isolated from control and cKO brains, without DTT in the sample buffer. PLP monomers and dimers are indicated with arrows. Quantification of normalized PLP-dimer levels between the two groups. Data information: Data are shown as mean ± SEM. n=3-4 animals per genotype; Student’s t-test was used. *p < 0.05. **p < 0.01.

First and foremost, we aimed to evaluate whether the main protein constituents of myelin, namely PLP and MBP, are altered in the mutants. Noticeably, we found that PLP protein levels were increased 2,5-fold as shown in the Western blot analysis of the 6mo cKO mice, while MBP levels did not show any difference (Figure 2 A, B). These data indicate that the autophagic machinery could be essential for the degradation of the PLP protein, since its dysregulation leads to accumulation of PLP. Since increased levels of PLP protein could also arise from a transcriptional dysregulation of the gene, we performed quantitative real-time PCR on brain extracts from 6mo control and cKO animals and measured the mRNA levels of plp and mbp genes, which came in accordance with the protein levels. In other words, plp mRNA was increased 1,5-fold in cKO compared to controls, while there was no difference in mbp mRNA levels between the two groups (Figure 2C). In the literature, increased copies in PLP1 gene and subsequently in PLP protein lead to an X-linked inherited demyelinating disorder, that of Pelizaeus-Merzbacher disease (PMD). Furthermore, it is known that mutant forms of PLP identified in patients of PMD, can form homo-oligomers and this formation may contribute to the pathophysiology of the disease process (Dhaunchak & Nave, 2007). To test this hypothesis, we isolated “native membrane extracts” (NME) of 6mo control and cKO forebrains in order to examine the possibility that the excess PLP protein in the cKO forms excessive disulfide-linked dimers of PLP. Western blot analysis of NME for PLP revealed no difference between control and cKO (Figure 2D).

We subsequently asked whether plp accumulation could be toxic to OL populations, as it occurs in PMD models (Skoff, 1995; Simons, 2002). To this end, and in order to study the influence of autophagy in an already myelin-mature system, we performed immunohistochemistry analysis against CC1 (mature OLs) and PDGFRa (oligodendrocyte progenitor cells, OPCs) in coronal sections of rostral corpus callosum of 6mo control and cKO mice (Figure S2A). No differences in the numbers of the different subpopulations were detected between the two groups (Figure S2B). The dispensable role of ATG5 in cell survival was further confirmed by the absence of cleaved caspase-3 activity in the region of the corpus callosum, a region rich in somata of OLs (Figure S2C). These results indicate that loss of autophagy in mature OLs does not cause their death, for at least 3 months following ATG5 deletion.

### Myelin proteins are degraded through autophagy

We set the hypothesis that autophagy may serve for the degradation of myelin proteins which consist cargoes and through the ablation of the mechanism in adult CNS, PLP protein accumulates *in vivo*. First, we aimed to validate the direct interaction of myelin proteins with autophagic vesicles. To this end, initially we performed primary OL cultures and verified the colocalization of the two most abundant protein components of CNS myelin, namely PLP and MBP, with the autophagosomal marker LC3-II (Figure 3A). To further validate that there is an interaction of these proteins with LC3-II, *in vivo* this time, we performed immunoprecipitation experiments using forebrain lysates from the GFP-LC3 transgenic line (Figure 3B) as well as an unbiased approach including autophagosome purification (Kallergi *et al*, 2022). We found that both PLP and MBP proteins are co-immunoprecipitated with fused LC3. Moreover, autophagic vesicles were isolated from adult wild type mouse forebrains and further assessed to a Proteinase K (PK) digestion step, in order to distinguish between the autophagosome cargo and the proteins attached to the outer membrane of the vesicle. Protein cargoes tend to be protected from the addition of PK, unless Triton-X is present. As shown in Figure 3C both proteins were protected in case of PK treatment and only digested in the presence of the Triton X-100 detergent, suggesting that they consist protein cargoes in autophagosomes.

**Figure 3:**
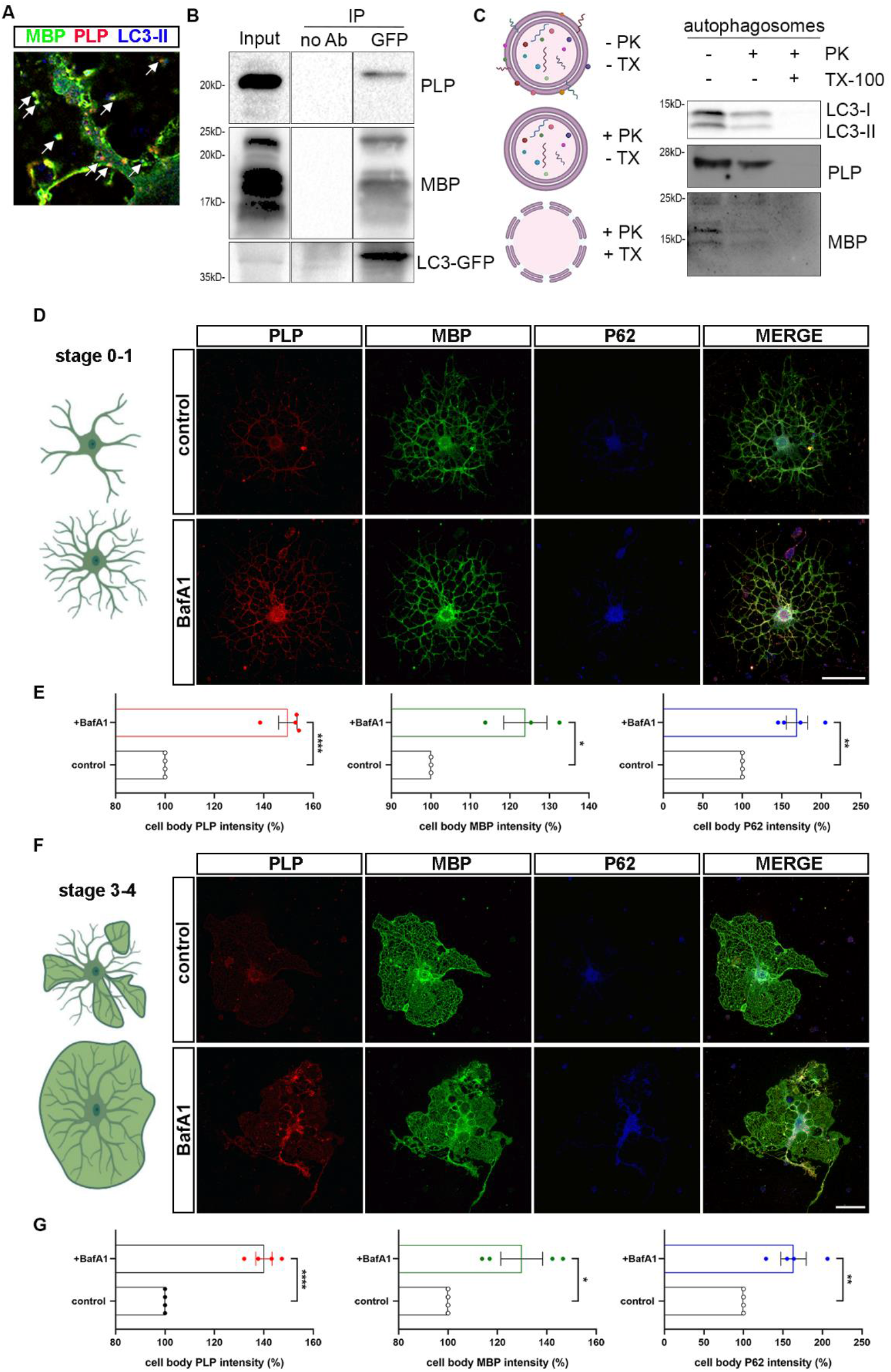
Myelin proteins PLP and MBP are detected in the autophagosomes and constitute autophagic cargoes. A. Confocal images of DIV2 primary OLs, immunostained for MBP (green), PLP (red), LC3-II (blue). White arrows indicate the colocalized puncta. B. Immunoprecipitation with antibodies against GFP in forebrain lysates from adult GFP-LC3 mice identifies MBP and PLP as interactors of LC3. C. Shematic of the proteinase K protection assay and western blot analysis of isolated autophagosomes after proteinase K assay. Triton X-100 (TX-100) is used as a negative control. Autophagic marker LC3-II was protected from PK digestion, unless TX-100 was present. Both PLP and MBP seem to be protected from PK treatment. D, F. Confocal images of DIV2 primary OLs, stage 0-1(A) and stage 3-4 (C), immunostained with antibodies against PLP (red), MBP (green) and P62 (blue). Cells were either vehicle treated (control), or treated for 4 hr with 10nM BafilomycinA1 (BafA1). E, G. Graph showing quantification of the normalized intensity levels of the above-mentioned proteins. Data information: Data are shown as mean ± SEM. Student’s t-test was used; n=4 independent experiments. *p < 0.05, **p≤ 0.01, ****p≤ 0.0001. Scale bars: 30 μm.

Additionally, to test that these two structural proteins consist cargo of the autophagosome and are not externally associated with the vesicle, we blocked the final step of the autophagic pathway, therefore preventing the lysis of the constituents of the autophagic vesicles. For this reason, we treated cultured primary OLs with Bafilomycin A1 (BafA1), an inhibitor of lysosome acidification. In order for the analysis to be more reliable, we subdivided our OLs in two different groups, namely stage 0-1 (OLs without myelin membranes, Figure 3D) and stage 3-4 (OLs with myelin membranes, Figure 3F), since there are significant differences in the signal intensity of proteins between those groups. Noticeably, BafA1-treated cells presented increased levels of p62 signal, a well-known substrate of autophagy, in both groups of OLs (Figure 3E, G). In parallel, we observed that beyond p62, PLP and MBP were also accumulated in the soma of OLs in case of BafA1 treatment in both stage 0-1 and stage 3-4 groups, indicating that these proteins use the autophagic machinery for their degradation (Figure 3E, G).

### Autophagy in oligodendrocytes is an essential mechanism for the maintenance of CNS myelin and its depletion causes behavioral deficits

The question that arose next was whether autophagy plays a role, in addition to the PLP homeostasis, also in myelin maintenance in the mature CNS. In order to test this hypothesis, we performed electron microscopic (EM) analysis of 6mo control and cKO optic nerves (Figure 4). This analysis revealed that in mutant mice there were significantly increased numbers of axons with decompacted myelin, as well increased numbers of degenerating axons (Figure 4C). Previous work on PMD, has established that a low-level increase of PLP dosage leads to late-on axonal degeneration (Anderson *et al*, 1998; Ip *et al*, 2006). Furthermore, quantification showed that there was no difference in the number of unmyelinated axons between the two groups. The lack of differences in the number of unmyelinated axons was expected since the ablation of autophagy in these mice is accomplished after myelination is complete. Additionally, the g-ratio analysis revealed that autophagy-deficient axons had smaller g-ratios, meaning thicker or extended myelin sheaths compared to controls (Figure 4D). This increase in myelin thickness could partly reflect the loosening of myelin lamellae or a thicker myelin sheath per se. Finally, analysis of axon diameters showed that there was a small, but statistically significant, redistribution of the percentages of axon caliber diameters, with the 6mo cKO axons presenting a loss of large caliber axons and increase in the pool of small caliber axons (Figure 4E). The above results, indicate that large caliber axons are more vulnerable in case of autophagic dysfunction and due to PLP accumulation. Larger diameter axons are needed to achieve greater conduction velocity and thus shorter conduction times (Rushton, 1951; Waxman & Bennett, 1972; Koch, 1999). Furthermore, it has been proposed that large caliber axons can support larger terminal arbors and more active zones that transfer information synaptically at higher rates (Perge *et al*, 2009, 2012). The large caliber axons are also positively associated with thicker myelin sheaths (Schroder *et al*, 1978), a characteristic that could render them more susceptible to the decompaction and loosening of myelin lamellae, observed in the 6mo cKO mice. Similar myelin abnormalities were revealed in other CNS tracts in our transgenic mice, such as the corpus callosum (data not shown).

**Figure 4:**
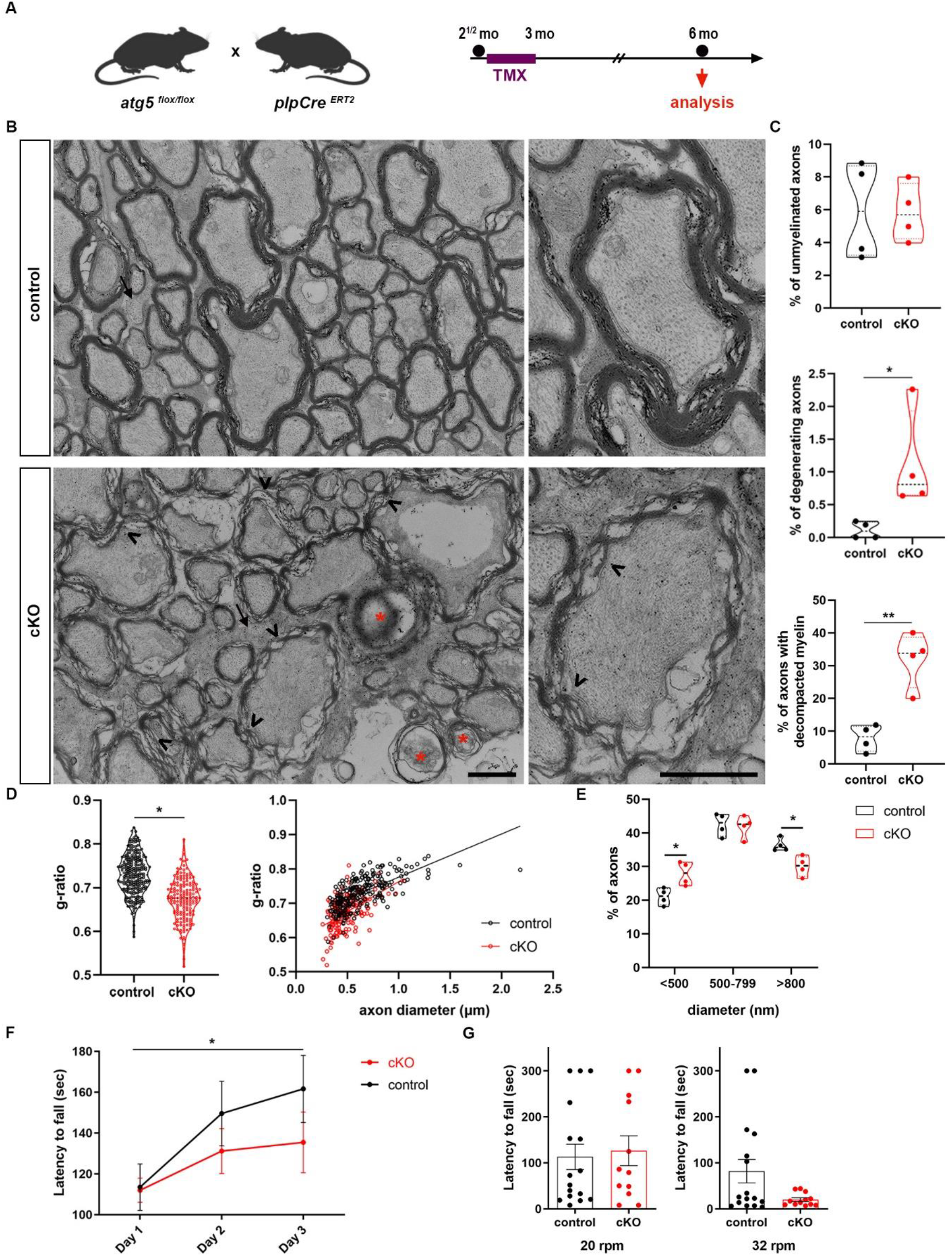
autophagy in OLs is an essential mechanism for the maintenance of CNS myelin. A. Schematic illustration of the experimental protocol used for tamoxifen (TMX) induction and analysis in the *plpCre*^*ERT2*^; *atg5*^*F/F*^ mice. All animals were injected i.p. with 1 mg of TMX per day at the age of 2.5 months for 10 days with two days break in between. The analysis was performed 3 months later (at 6 mo). B. Ultrastructural analysis of optic nerves from *plpCre*^*ERT2*^*-; atg5*^*F/F*^ (control) and *plpCre*^*ERT2*^*+; atg5*^*F/F*^ (cKO) mice. Representative electron micrographs of cross sections of control and cKO optic nerves, demonstrating the presence of degenerating axons (indicated by red stars), decompacted myelin sheaths (arrowheads) and unmyelinated axons (arrows). C. Quantification of axonal degeneration, decompacted myelin sheaths and unmyelinated axons in control and cKO mice. Violin plots presenting individual values (450-500 axons measured for each mouse, unpaired two-tailed t-test. N= 4 animals per genotype). D. Myelin sheath thickness was assessed using g-ratio measurement of EM images from control and cKO animals. Plot of g-ratio values per animal group (each point represents an axon) and g-ratio distribution to different axonal diameters of the two groups (linear regression, 170 axons measured for cKO group and 260 for the control, n= 4 animals per genotype). Scale bar: 1 μm. E. Quantitation of the percentage of axons with small (<500nm), medium (500-799nm) and large (>800nm) caliber in control and cKO optic nerves. (250 axons measured for each mouse, unpaired two-tailed t-test. N= 4 animals per genotype. *p < 0.05, **p < 0.01, n.s. = non-significant. F. Performance of mice in the rotarod task. Line graph showing latency to fall during training (3 days). N= 16 for control; n=12 for cKO. G. Line graph showing latency to fall during the testing phase in the rotarod task. Control, n= 16; cKO, n=12. Data information: Bar graphs depict mean ± SEM.

Our findings above raise the question whether the blockage of autophagy in OLs, in addition to defects in myelin integrity, also leads to functional impairments. To elucidate this hypothesis, we examined behaviors such as motor learning and motor coordination, which are associated with myelin integrity, in the brain of 6mo cKO and control mice. To test whether motor learning is altered in case of defective autophagy, mice had to perform an accelerating rotarod test for three consecutive days (Costa *et al*, 2004). In this task, 6mo control mice showed a continuous increase in the time required until they fell within the 3-day trial period, meaning they significantly improved their performance over time. On the other hand, age-matched cKO littermates did not show any statistically important improvement in their performance over time, indicating that they were not able to learn this skilled motor task (Figure 4F). Following the 3-day accelerating trial (Figure 4F), mice performed the motor coordination testing phase of the task, in which the latency to fall in a rod rotating in a constant speed of 20rpm and 32rpm respectively is tested for the next two days (Figure 4G). A trend of lower latency to fall was detected in cKO mice, only in the higher speed task (Fig 4G), suggesting that 6mo cKO mice have a mild dysfunction in their motor coordination.

Overall, our *in vivo* analysis proposes an important role for autophagy as a homeostatic mechanism for the maintenance of proper PLP levels, thus the accumulation of this protein in oligodendrocytes is accompanied by myelin and axon morphological defects as well as defects in cognitive behavior in mice.

## Discussion

Our findings indicate that autophagy is an essential mechanism for OL maturation, since both the pharmacological and genetic inhibition of autophagic vesicle formation led to a final maturation defect in primary OL cultures (fewer stage 4 cells). Furthermore, we showed with biochemical and immunostaining techniques that the structural myelin proteins PLP and MBP use this pathway for their degradation. Specifically, both these proteins are detected in autophagosomes, although only PLP accumulates *in vivo* in case of autophagic blockage. This may be due to the fact that PLP is more abundant in CNS myelin [it consists ∼50% of CNS myelin protein, while MBP about 30% (Boggs, 2006)]. In addition, MBP may not be accumulated because it uses a different mechanism of expression at the site of interest. The majority of MBP protein in translated locally under the OL membrane in contrast to PLP. No excess MBP is detected under the myelin membrane since its translation responds to external stimuli (i.e. axon stimulation). Thus, cargo MBP for autophagy accounts only for a small portion of the protein, possibly the cytoplasmic and nuclear form which consist of only a small portion of the total and therefore, small differences may not be detected. Alternatively, MBP may be degraded via other mechanisms such as the proteasome that directly takes part in the specific degradation of MBP (Bacheva *et al*, 2009). Moreover, in the case of PLP, both protein and mRNA levels were significantly increased in case of autophagic depletion in oligodendrocytes. These increased transcriptional levels could imply the possible role of autophagy not only in protein degradation, as is well characterized, but also in mRNA degradation, a pathway that has only very recently been described in yeast (Makino *et al*, 2021). Another potential explanation for this increase could be that the accumulated PLP protein levels are not functional. It is possible to envisage that the cell does not recognize this nonfunctional PLP, leading to the production of more mRNA due to compensation. Increased plp mRNA expression does not automatically correspond to increased functional PLP protein, as has already been described in the literature in the demyelinating model of PMD (Mayer *et al*, 2011).

Moreover, in this study we ablated the core autophagic gene *atg5* in OLs after myelination is completed (at 2.5 mo) and analyzed these animals at 6mo. Recent work, targeting OPCs, has shown reduced and defective myelination in case of an early OPC-specific deletion of atg5 autophagy (Bankston *et al*, 2019). In our study, in case of autophagic ablation, we observe significantly increased numbers of axons with decompacted myelin and axonal degeneration. The differences observed in the severity of the phenotype between the two studies may be explained first and foremost due to the fact that Bankston *et al* ablate autophagy in the stage of the progenitors and at an early developmental stage, while in our case, by ablating autophagy in adulthood, we focus in myelin maintenance, not its formation. Secondly, the severity of the phenotype stated in Bankston *et al* may rely to some degree to the transgenic line they use, which targets not only OPCs, but also pericytes, cells that are implicated in CNS myelination and whose dysfunction has been recently related to white matter changes (Montagne *et al*, 2018; Azevedo *et al*, 2018). Furthermore, our electron microscopy analysis revealed a loss of large caliber axons. According to the literature, a shift towards larger axon calibers is observed in ageing optic nerves (Stahon *et al*, 2016), as well as in pathological cases, such as during chronic secondary degeneration in rat optic nerves (Payne *et al*, 2012). Moreover, 6mo cKO animals present behavioral deficits such as decreased motor skill learning, an ability that is directly correlated to active central myelination (McKenzie *et al*, 2014; Kato *et al*, 2020). In general, one could notice that this phenotype resembles an “early-ageing” phenotype. In the literature, ageing has been associated with impairments in learning as well as increased degeneration and decompaction of myelin sheaths (Sandell & Peters, 2001, Peters 2009). In parallel, there is evidence showing that autophagy decreases during ageing, while myelin proteins are increased (Aman *et al*, 2021; de la Fuente *et al*, 2020).

Overall, we propose two different hypotheses supported by our results. Firstly, it is possible that all the abnormalities observed in case of ablated autophagy in OLs are a result of PLP accumulation, similar to the case of PMD. In this disease, patients with PLP1 duplication present segmental demyelination along axons, abnormally thick myelin sheaths and degeneration of axons (Laukka *et al*, 2016), a phenotype that extensively resembles our case. Furthermore, it is noteworthy to mention that in the PMD mouse model, an increased number of autophagosomes has been described which would corroborate our findings (Karim *et al*, 2010). An alternative explanation of this phenotype could be a result of accumulation of damaged or excess myelin membranes per se, meaning that autophagy is responsible for degradation of CNS myelin. This hypothesis is supported by the fact that two major myelin proteins (PLP and MBP) are degraded through autophagy, leading to the idea that CNS myelin itself could use this machinery for its degradation. Our existing knowledge documents that the elimination of aberrantly deposited myelin -in case of trauma or during development-is accomplished through phagocytosis via microglia as well as endocytosis via astrocytes (Huges & Appel, 2020; Ponath *et al*, 2017; Zhou *et al*, 2019). Thus, our hypothesis is that, in addition to these mechanisms, autophagy in Ols could also clear CNS myelin deposits under steady state conditions, an assumption that requires further investigation.

## Materials and methods

### Mouse lines

The animal protocols of this study were approved by the Animal Ethics Committee of the Foundation for Research and Technology Hellas (FORTH). All animals were kept at temperature-controlled conditions on a 12 h light/dark cycle, fed by standard chow diet and water ad libitum provided within the animal facility of the Institute of Molecular Biology and Biotechnology (IMBB) - Foundation for Research and Technology Hellas (FORTH) (license nos. EL91-BIObr-01 and ?L91-BIOexp-02). All animal experiments complied with the ARRIVE and NC3Rs guidelines to improve laboratory animal welfare and conformed with all regulations and standards outlined in the Presidential Decree 56/ 30.04.2013 (Greek Law) in accordance with the EU directives and regulations (2010/63/EU and L 276/33/20.10.2010) and to the U.K. Animals (Scientific Procedures) Act, 1986 and associated guidelines, equivalent to NIH standards. The ages of the mice are described in each experiment and only males are used in this study. The animals used were of C57BL/6 genetic background. *Atg5* ^*F/F*^ (Hara *et al*, 2006;), *plpCre*^*ERT2*^ (Leone *et al*, 2003; a generous gift from Dr. Shuter, Zurich), *mT/mG* (Muzumdar *et al*, 2007; a generous gift from Dr. Talianidis, IMBB-FORTH) and *nestin-Cre (*Hara *et al*, 2006) mouse lines were used. In *plp-Cre*^*ERT2*^; *atg5* ^*F/F*^ and *mT/mG; plpCre*^*ERT2*^ progeny, tamoxifen (Sigma-Aldrich, T5648) was administered by intraperitoneal injections at a dose of 1mg/mouse/day for ten days with a two-day break in between, as previously described (Leone *et al*, 2003), at 2.5 months (mo) and mice were sacrificed at 6 months (6mo).

### Assessment of mouse motor phenotypes

#### Rotarod test

To assess the acquisition of motor behavior in mice, a rotarod apparatus with automatic timers and falling sensors was used (MK-660D, Muromachi-Kikai, Tokyo, Japan). The training was performed on the apparatus and consisted of four trials per day (with 15 min rests between trials), for 5 consecutive days. In the first three days, mice were placed on the rotating rod at 4 rpm and gradually the speed was increased to 40 rpm. Each trial lasted until the mouse fell from the rod or for a maximum of 300s. In the fourth and fifth day, the task consisted of two consecutive sessions of three trials each (maximum duration, 300s): the first session at a constant speed of 20 rpm, and the second one at 32 rpm. For the analysis the time until the mice dropped from the rod for each of the training days as well as for each testing speed was measured (Savvaki *et al*, 2010; Lee *et al*, 2016).

#### Statistical Analysis

Statistical analyses regarding the behavioral tests were performed using SPSS version 25 (SPSS Inc., Chicago, Illinois). Between-group differences in the tasks in control and cKO mice were examined either with parametric or non-parametric analyses according to normality of the distribution, as examined with the Kolmogorov–Smirnov test. For the rotarod task the effect of genotype on the latency before falling off during training, was assessed using paired t-test. For the 4th (20rpm) and 5th day (32rpm) of the task (testing phase) the effect of genotype on the latency was assessed using one-way ANOVA.

#### Immunohistochemistry and immunocytochemistry

For immunohistochemistry, brains and optic nerves were harvested after transcardial perfusion with 4% paraformaldehyde (PFA) in 0.1 M phosphate buffer saline (PBS). Tissues were then post-fixed in the same fixative for 30 min at 4°C, cryo-protected overnight in 30% sucrose in 0.1 M PBS, and embedded in 7.5% gelatin/ 15% sucrose gel. Cryosections of 10 μm-thick for optic nerves and 15 μm -thick sagittal or coronal for brains were obtained and mounted on Superfrost Plus microscope slides (O. Kindler), post-fixed in ice-cold acetone for 10 min, blocked in 5% BSA (Sigma-Aldrich) in 0.1 M PBS for 1h at RT, and incubated with primary (overnight at 4°C) and secondary (2h at RT) antibodies in 5% BSA, 0.5% Triton-X in 0.1 M PBS.

For immunocytochemistry, cells were fixed with 4% PFA for 15 min at RT, blocked/ permeabilized for 30 min with 1%BSA, 0.1% Triton-X in 0.1 M PBS and incubated with the primary antibody and secondary antibodies in 1%BSA, 0.1% Triton-X in 0.1 M PBS. Sections or coverslips were mounted using MOWIOL Reagent (Merk-Millipore, Burlington, MA) and image acquisition was performed in a confocal microscope (TCS SP8, LEICA DMI-8).

The following primary antibodies were used: anti-PLP (1:1000, rabbit, Abcam Cat# ab28486), anti-MBP (1:200, rat, Serotec), anti-adenomatous polyposis coli clone CC1 (APC/CC-1) (1:100, mouse, Millipore Cat# OP80) anti-platelet derived growth factor receptor alpha (PDGFRa) (1:100, rat, Millipore Cat# CBL1366), anti-GFP (1:2000, rat, Nacalai Tesque, Cat# 04404-26), anti-p62 (1:500; rabbit, MBL, Cat# PM045), anti-LC3b (1:1000; rabbit, Sigma, Cat# L7543).

Fluorochrome labeled secondary antibodies Alexa Fluor 488, 555 and 633 (Invitrogen, Carlsbad, CA, 1:800) were also used. DAPI (Dako, Hamburg, Germany) was finally used for the visualization of the nuclei.

### Electron microscopy

#### Tissue preparation

For electron microscopy, mice were perfused with 2.5% glutaraldehyde in 0.1 M phosphate buffer (PB), pH 7.25. Entire dissected optic nerves were placed in the primary fixative overnight at 4°C and then the preparation of the samples for the analysis was done similarly to Pasquettaz *et al*, 2020. Briefly, the samples were extensively washed in PB 0.1 M and incubated in 2% (wt/vol) osmium tetroxide and 1.5% (wt/vol) K4[Fe(CN)6] in 100 mM PB buffer for 1h on ice. After an extensive wash with water, samples were incubated for 1h in 1% (wt/vol) tannic acid in 100 mM PB buffer, followed by 1% (wt/vol) uranyl acetate for 2h at ambient temperature. Finally, samples were dehydrated at ambient temperature in gradual ethanol cycles and infiltrated with a mix of ethanol and Epon-Araldite mix (EMS). After several cycles of 100% Epon-Araldite incubations, samples were flat embedded and polymerized for 24h at 60°C (Kolotuev, 2014). The experimental procedure was performed simultaneously in control and cKO samples, in order to avoid the possible implication of artifacts arising from fixation or embedding.

#### Electron microscopy

Polymerized flat blocks were trimmed using a 90° diamond trim tool (Diatome, Biel, Switzerland). The arrays of 70 nm sections were obtained using a 35° diamond knife (Diatome, Biel, Switzerland) mounted on Leica UC6 microtome (Leica, Vienna). The orientation for the optic nerves was established perpendicular to its length, a cross-section about 1 mm from the optic nerve head. For sectioning, samples were carefully oriented to obtain a perpendicular plane of the optical nerve. In the case of brain slices, the search was targeted to the corpus callosum based on the overall morphology of the processed slice (Kolotuev, 2014, Burel *et al*, 2018). Sections were collected on polyetherimide-coated carbon slot grids.

TEM samples were analyzed with an FEI CM100 electron microscope (Thermo Fischer Scientific) at 80kV, equipped with a TVIPS camera, piloted by the EMTVIPS program. Images were collected either as single frames or stitched mosaic panels to cover more extensive sample regions.

The multiple tile images were stitched with the IMOD software package (Kremer *et al*, 1996). Data were processed and analyzed using Fiji, IMOD 3dmod, and Photoshop programs.

#### Western blot analysis and immunoprecipitation

Tissues were collected and stored at -80^°^C until their homogenization. The number of animals for each experiment is indicated in the results section. Briefly, tissues were lysed by sonication in RIPA buffer (500 mM Tris-HCl pH 7.2, 1MNaCl, EDTA, Triton 100-X, Na-deoxycholate, 10% SDS), supplemented with protease inhibitors (Sigma) and 1 mM dithiothreitol (DTT), and placed for 20 min on ice, followed by 20 min centrifugation at 14,000 rpm. Samples were separated on a 10% or 15% polyacrylamide gel and transferred to a nitrocellulose membrane (Millipore). After blocking for 1 hr at room temperature in 5% BSA (sigma), membranes were incubated in the primary antibodies overnight at 4°C. The following antibodies were used for Western blot (WB) analysis: anti-PLP (1:1000, rabbit, Abcam Cat# ab28486), anti-MBP (1:200, rat, Serotec), anti-p62 (1:1000; rabbit, MBL, Cat# PM045), anti-LC3b (1:1000; rabbit, Sigma, Cat# L7543).

After three 5 min washes in TPBS (100 mM Na2HPO4, 100mM NaH2PO4, 0.5N NaCl, 0.1% Tween-20), membranes were incubated for 1h at room temperature in corresponding secondary horseradish peroxidase-conjugated antibodies (Abcam). Blots were developed by chemiluminescence (Immobilon Classico Western HRP substrate, Merck, WBLUC0500) according to the manufacturer’s instructions. Quantification of band intensity was assessed with the Fiji/ImageJ Gel Analyzer plugin.

For co-immunoprecipitation, SureBeads Protein G Magnetic Beads (Biorad, Cat# 161-4023) were used, following the protocol suggested by the manufacturing company. 3 μl of a polyclonal antibody for GFP, was used for each experiment and 300–500 μl of protein lysate (1–2 mg total protein from cerebellum lysates). Immunopurified material was then used for western blot.

#### Biochemical purification of mouse autophagosomes (AVs) and Proteinase K protection assay

Ten cortices and hippocampi of adult (postnatal day 60, P60) C57BL/6J male mice were used to purify brain AVs, as described previously (Nikoletopoulou *et al*, 2017). Briefly, the aforementioned brain areas were homogenized in 10% sucrose, 10 mM Hepes and 1 mM EDTA (pH 7.3) by 20 strokes using a Dounce glass homogenizer. The resulting homogenate was sequentially dilluted with half volume of homogenization buffer (HB) (250 mM sucrose, 10 mM Hepes and 1 mM EDTA, pH 7.3) containing 1.5 mM glycyl-L-phenylalanine 2-napthylamide (GPN). The resulting material was then incubated at 37°C for 7 min, and centrifuged at 2000g for 2 min at 4°C. The nuclear pellet was discarded and the post-nuclear supernatant was loaded on discontinuous Nycodenz gradients for centrifugation at 141.000g for 1h at 4°C, to remove the cytosolic, mitochondrial and peroxisomal fraction. The isolated layer of material, which contained both autophagosomes and endoplasmic reticulum, was further diluted with an equal volume of HB buffer and overlaid on Nycodenz-Percoll gradients. After centrifugation at 72000g for 30 min at 4°C to remove the Percoll silica particles, the resulting interface that contained the AVs was diluted with 0.7 volumes of 60% Buffered Optiprep overlaid by 30% Buffered Optiprep and HB buffer. The gradients were then centrifuged at 71000g for 30min at 4°C. The collected AVs were diluted in three volumes of HB buffer and the concentration of the purified isolated AVs was then measured by BCA, following the manufacturer’s instructions. For the Proteinase K (PK) protection assay, AVs were treated with PK (20ng/μl) on ice for 20 min, in the presence or absence of 1% Triton X-100, and then PK was inactivated using 4 mM of PMSF for 10 min on ice. The AVs were then centrifuged at 16.900g for 10 min at 4°C, and the autophagosomal pellets were resuspended in Laemmli buffer and boiled at 95°C for 5min for immunoblotting analysis. The purity of autophagosomes isolated out of this protocol and the absence of contamination by other organelles is already established (Kallergi *et al*, 2022).

#### Real time PCR

Total RNA was prepared from cortical samples from control and cKO animals (3 animals/group) by using RNAiso-plus kit (Takara) and according to the manufacturer’s instructions. cDNAs were synthesized through reverse transcription from the total RNA according to the protocol of Affinity Script Multiple Temperature cDNA synthesis kit (Agilent Technologies, Santa Clara, CA). Expression levels of genes encoding plp (forward primer: 50-TCAGTCTATTGCCTTCCCTA-30, reverse primer: 50-AGCATTCCATGGGAGAACAC-30), mbp (Pernet *et al*, 2008; forward primer: 50-CACACACGAGAACTACCCA-30, reverse primer: 50-GGTGTTCGAGGTGTCACAA-30), were examined by real-time PCR analysis using a StepOnePlus real-time PCR system (Applied Biosystems, Life Technologies, Thermo Fisher Scientific Inc., Waltham, MA). Gapdh was used as the internal control (forward primer: 5’-ATTGTCAGCAATGCATCCTG-3’, reverse primer: 5’-ATGGAC TGTGGTCATGAGCC-3’). PCR runs were performed for each sample in triplicates and the expression levels for each gene were normalized according to the internal control.

#### Primary OL cultures

Primary OL cultures were obtained from postnatal day 2 (P2) mouse cortices, as previously described (Zoupi *et al*, 2017). OLs were seeded at an initial density of 70,000 cells per well in 24-well plates containing 13mm glass coverslips or at an initial density of 35,000 cells per well in 48-well plates containing 9mm glass coverslips. All plates were previously coated overnight with poly-D-lysine (Sigma–Aldrich, A-003-E). OLs were cultured in DMEM (High glucose-pyruvate, ThermoFisher Scientific, Cat# 61965-026), supplemented with 1% N2 (ThermoFisher Scientific, Cat#17502), 1 μM D-biotin (Sigma-Aldrich, Cat# B4501), 1% BSA fatty acid-free (Sigma-Aldrich), 5 μg/ml N-acetylcycteine (Sigma-Aldrich, Cat# A8199), 1% penicillin-streptomycin (ThermoFisher Scientific, #15070063) and 40 ng/ml T3 (Sigma-Aldrich, Cat# T6397) to allow the differentiation of OPCs toward mature OLs. For experiments involving SBI-0206965 treatment, SBI-0206965 (Sigma-Aldrich #SML1540, diluted in 0.1% DMSO) was used at a concentration of 1uM and replenished every 48 h until DIV5, when cells were fixed. Bafilomycin A1 (Sigma-Aldrich #B1793, diluted in 0.1% DMSO) was added in the medium for 4h (10nM) before fixation. For both treatments, control OLs were treated with the vehicle at the same concentration (0.1% DMSO) and for the same time duration.

#### Cell viability assay

For the cell viability assay, OLs were seeded at an initial density of 10,000 cells per well in 94-well plates for 3 days. Then cells were treated with SBI-0206965 (1uM) for 24h or 48 h, and cell viability was evaluated via 3-(4, 5-dimethylthiazol-2-yl)-2, 5-diphenyltetrazolium bromide (MTT) assay (Molecular Probes, Cat# V-13154). The analysis was conducted in triplicate and three independent experiments were used. Cell viabilities were defined relative to control cells (vehicle treated, considered to be 100%). The absorbance was read at 540 nm using the Infinite® 200 PRO NanoQuant microplate reader (Tecan, Research Triangle Park, NC, USA).

#### Statistical analysis

In all experiments, data were expressed as mean ± SEM. Parametric tests including two-tailed, unpaired t test and one-way analysis of variance (ANOVA), followed by Tukey post-hoc test for multiple comparison procedures were performed where values follow a Gaussian (normal) distribution. In the case of non-normally distributed values, non-parametric Mann–Whitney test or Kruskal– Wallis test with Dunn’s post hoc tests for multiple comparisons were used. Statistical analysis, in all the experiments except from the behavioral tests, was performed using GraphPad Prism 8 software (GraphPad Software, San Diego, CA). P-values 0< 0.05 were considered statistically significant.

## Data availability

Data sharing is not applicable to this article as no datasets were generated or analyzed during the current study.

Expanded View for this article is available online.

## Acknowledgements

We are thankful to Prof. A. Stamatakis and Prof. K. Sidiropoulou for their help and advice in the behavioral analysis. We would like to acknowledge funding from the Hellenic Foundation for Research and Innovation (HFRI grant agreement 1676), the National Multiple Sclerosis Society (NMSS, pilot Research Grant, PP-1809-32556), Fondation Santé, the Company of Biologists (Travelling Fellowship to N.K.), the Boehringer Ingelheim Fonds (travel grant to N.K.) and the Manassaki graduate fellowships of the University of Crete (to N.K.). Furthermore, we would like to state that cartoons in Figures 3c-f, 4a and S1a were created with BioRender.com.

## Supplementary figure legends

**Figure S1:**
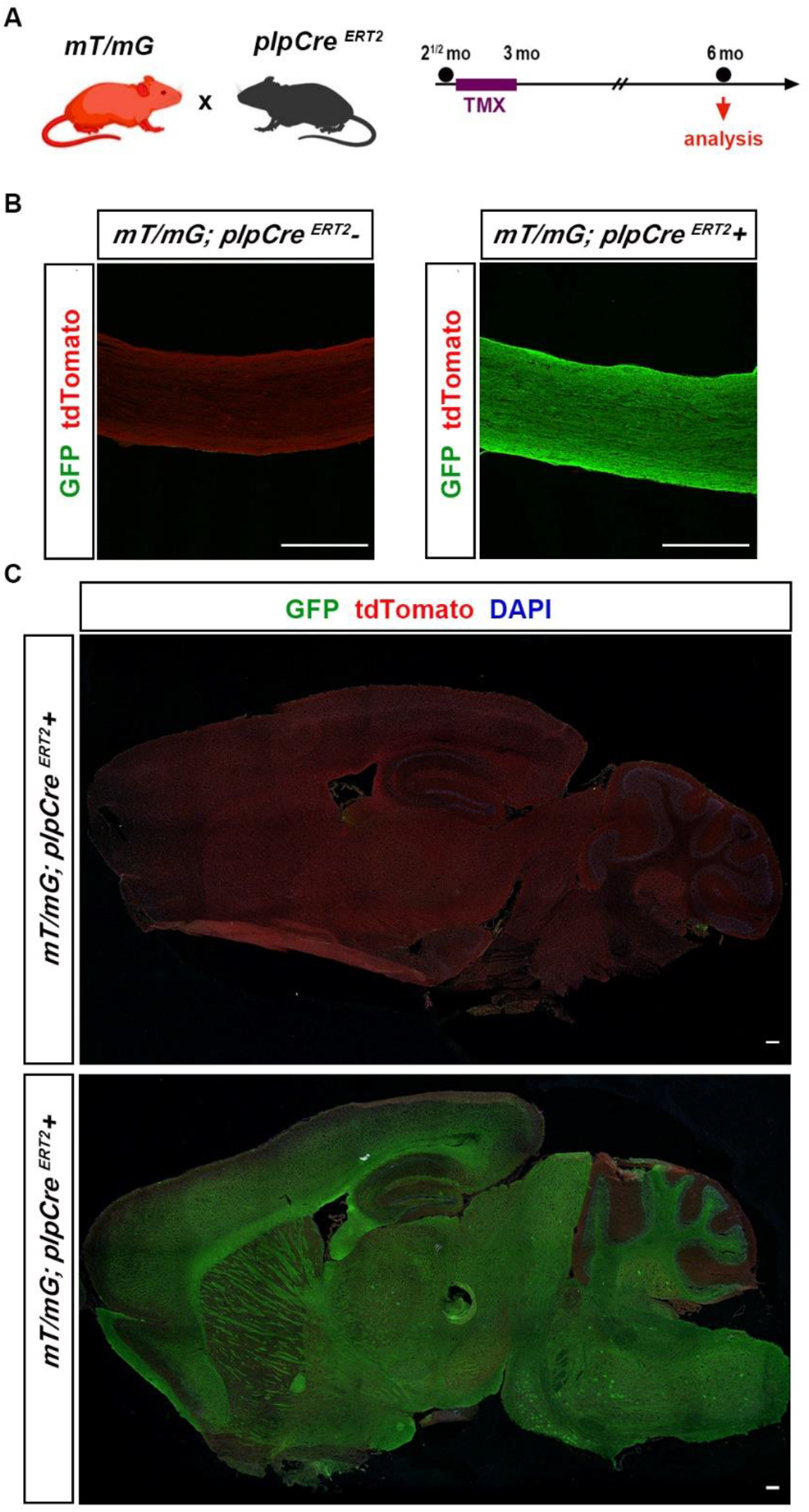
Recombination efficiency of tamoxifen inducible *plpCre*^*ERT2*^ line. A. Schematic illustration of the experimental protocol used for TMX induction and analysis in the *plpCre*^*ERT2*^; *atg5*^*F/F*^ mice. All animals were injected i.p. with 1 mg of TMX per day at the age of 2.5 months for 10 days with two days break in between. The analysis was performed 3 months later (at 6 months of age). B. Representative images of the percentage of GFP+ (green) recombinant cells in optic nerve cryosections of *mT/mG; plpCre*^*ERT2*^+ and *mT/mG; plpCre*^*ERT2*^*-* mice. Intense green fluorescent is detected only in plpCre+ optic nerves at both ages. C. Recombination efficiency in sagittal brain cryosections of 6mo *mT/mG; plpCre*^*ERT2*^*+* and *mT/mG*^*+/-*^; *plpCre* ^*ERT2*^-mice which have received 10 doses of 1mg of TMX at 2.5 mo. Confocal images of the percentage of GFP+ (green) recombinant cells (nuclei are stained with DAPI in blue) reveals green fluorescent in myelin tracts only in *plpCre+* transgenic mice. Data information: Scale bars: 200 μm.

**Figure S2:**
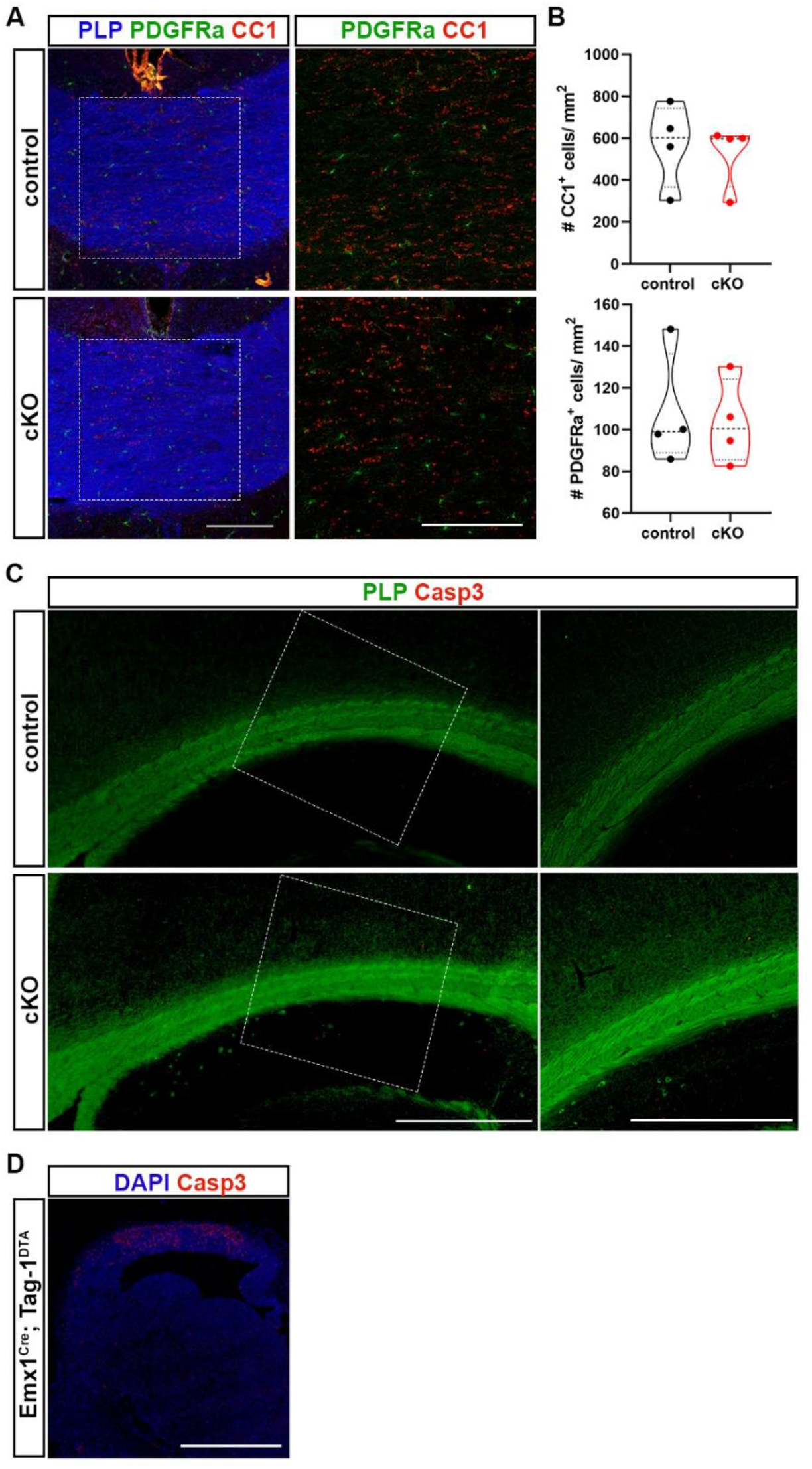
ATG5 is dispensable for OL survival. A. Immunohistochemical analysis of rostral corpus callosum (cc) cryosections from 6 mo transgenic and control mice. The area of cc is recognized with immunostaining against PLP (in blue). Two OL lineage markers were used: PDGFRa (in green) for OPCs and CC1 (in red) for mature OLs. B. Graphs showing the density of PDGFRa+ OPCs and CC1+ mature OLs in cc. C. Representative confocal images of sagittal cc sections from control and cKO mice immunostained for PLP (in green) and cleaved Caspase-3 (CASP3, in red). Rectangular boxes indicate areas magnified to the right. D. Representative confocal image of coronal sections from E13.5 Emx1^Cre^; Tag-1^DTA^ mice for cleaved Caspase-3 (CASP3, in red) and DAPI (in blue), used as a positive control of Casp3 staining. Data information: N=4 animals per genotype; Student’s t-test was used. Scale bars: 200μm (A), 500 μm (C, D).

